# Expanding the chemical space of synthetic cyclic peptides using a promiscuous macrocyclase from prenylagaramide biosynthesis

**DOI:** 10.1101/2020.03.17.996157

**Authors:** Snigdha Sarkar, Wenjia Gu, Eric W. Schmidt

**Affiliations:** Department of Medicinal Chemistry, University of Utah, Salt Lake City, Utah 84112, United States

## Abstract

Cyclic peptides are excellent drug candidates, placing macrocyclization reactions at the apex of drug development. PatG and related dual-action proteases from cyanobactin biosynthesis are responsible for cleaving off the C-terminal recognition sequence and macrocyclizing the substrate to provide cyclic peptides. This reaction has found use in the enzymatic synthesis of diverse macrocycles. However, these enzymes function best on substrates that terminate with the non-proteinogenic thiazole/thiazoline residue, complicating synthetic strategies. Here, we biochemically characterize a new class of PatG-like macrocyclases that natively use proline, obviating the necessity of additional chemical or biochemical steps. We experimentally define the biochemical steps involved in synthesizing the widespread prenylagaramide-like natural products, including macrocyclization and prenylation. Using saturation mutagenesis, we show that macrocyclase PagG and prenyltransferase PagF are highly promiscuous, producing a library of more than 100 cyclic peptides and their prenylated derivatives *in vitro*. By comparing our results to known cyanobactin macrocyclase enzymes, we catalog a series of enzymes that collectively should synthesize most small macrocycles. Collectively, these data reveal that, by selecting the right cyanobactin macrocyclase, a large array of enzymatically synthesized macrocycles are accessible.

## INTRODUCTION

Macrocycles are prevalent in drug development, with a growing number of macrocyclic peptides that are FDA approved or in clinical trials.^1–3^ However, the therapeutic scope of cyclic peptides remains underexploited. Synthetic macrocyclizing approaches include metathesis,^4–6^ click chemistry,^7–9^ the use of auxiliaries,^10–11^ and intein-mediated ligation,^12^ but despite many innovations limitations remain.^13^ Enzymes provide an additional tool to create libraries of cyclic peptides for drug discovery. Examples include subtiligase,^14–15^ cyanobactin macrocyclases,^16^ and cyclotide macrocyclases,^17–20^ each of which has been exploited in drug discovery programs. As with chemical synthesis, each enzyme has its limits, including restricted substrate tolerance, reaction condition requirements, and others. Here, we sought to remedy these limitations by exploring orthogonal cyanobactin macrocyclases for their biotechnological potential.

Cyanobactins are ribosomally synthesized and post-translationally modified peptides (RiPPs) found in cyanobacterial blooms.^21–22^ Most of these compounds are N-C circular, with macrocyclization catalyzed by a unique series of subtilisin-like serine proteases exemplified by PatG from *pat* pathway.^23–24^ The substrates for PatG are short precursor peptides containing the substrate for macrocyclization and a C-terminal recognition sequence (RS). A unique helix-turn-helix motif in PatG, the capping helix (CH), selectively recognizes the RS secures the substrate in the active site pocket.^25^ Subsequently, the PatG active-site serine proteolyzes the substrate, releasing the RS and leaving a covalently captured enzyme-substrate ester. In most serine proteases, a hydrolytic event then releases the substrate as a linear peptide.^12^ In PatG and relatives, instead of this canonical reaction, the covalent intermediate is captured by the substrate’s own N-terminus, leading to the synthesis of a cyclic peptide (Figure 1).^25^

**Figure 1.**
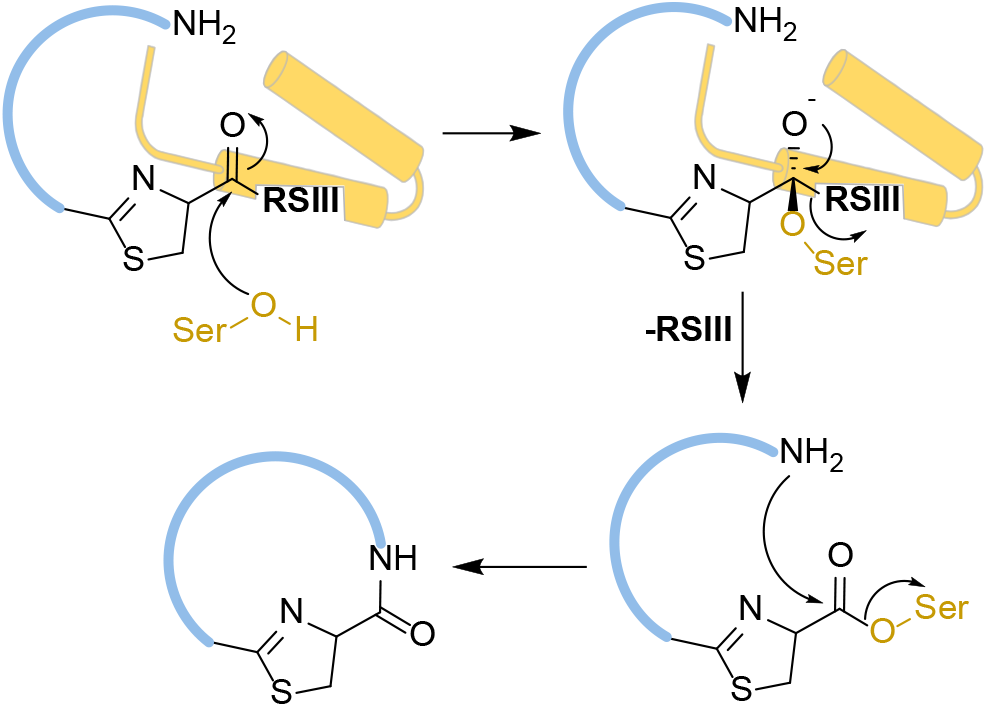
Proposed reaction mechanism of previously characterized cyanobactin macrocyclases, which prefer thiazoline residues. The recognition sequence ‘RSIII’ places the core peptide (blue) in the active site as a result of salt bridge interactions with capping helices (CH, yellow). Serine residue at the active site attacks the carbonyl carbon linking the core peptide and recognition sequence to form an acyl-enzyme intermediate. Finally, the N-terminus attacks the ester to produce the cyclic peptide. In this study, we biochemically characterize proline specific macrocyclases.

PatG and its homologs are a class of remarkably promiscuous catalyst, with studies *in vivo* and *in vitro* revealing a broad tolerance to accept natural and unnatural amino acids.^26–28^ Libraries encoding millions of products have been synthesized in *Escherichia coli* by TruG, a homolog of PatG from *tru* pathway.^29^ However, PatG has several limitations. It is a slow enzyme, at least under present conditions, and therefore disfavored substrates are impractical. PatG, and other macrocyclases that have been characterized so far, natively act on azolines. While proline and other heterocycles in artificial substrates is sometimes accepted by PatG, the reaction is usually much slower than it is with azoline.^30–31^ Generally, azolines are installed by a specific cyanobactin heterocyclase enzyme prior to macrocyclization.^32^ Although the heterocycle provides rigidity to the backbone to assist macrocyclization, this additional biochemical step increases complexity and compromises the yield for downstream applications such as library generation.

Over the years, there has been an increasing number of cyanobactin biosynthetic gene families, which afford diverse, broad-substrate enzymes for synthetic biology.^22, 33–34^ Among these is a group of cyclic cyanobactins that contain proline, rather than thiazoline, at the macrocyclization site. These pathways lack both azoline and the azoline-synthesizing heterocyclase enzyme, and thus might provide simpler macrocyclization catalysts. Unfortunately, no biochemical experiments for proline-specific cyclases have been reported, and in our hands, these enzymes have been challenging.

The first pathway for proline-selective cyanobactins was sequenced in 2009,^35^ and many more have since been identified, including the prenylagaramide (*pag*) pathway.^34^ Prenylagaramides, are a family of N-C cyclic, tyrosine O-prenylated peptides from *Planktothrix agardhii* cyanobacteria.^36^ Although *pag* has been cloned, only the PagF tyrosine O-prenyltransferase has been biochemically characterized.^37^ Here, we describe the characterization of PagG, a proline-selective macrocyclase that greatly expands accessible substrates for recombinant macrocyclases.

## RESULTS AND DISCUSSION

### PagG expression and *in vitro* activity

The *pagG* gene was codon-optimized and synthesized using the published sequence (GenBank accession number HQ655154).^34^ Despite attempting a wide variety of conditions, the full length PagG could not be produced in soluble form (Figures 2 and S1). The protein is comprised of an N-terminal protease/macrocyclase domain and a domain of unknown function (DUF) at the C-terminus.^25^ Previous experiments with PatG and TruG macrocyclases demonstrated that only the protease/macrocyclase domains were required for macrocyclization.^30^ The PagG macrocyclase domain (PagG^mac^) could be positively identified, expressed and purified, and this protein was used for all further experiments (Figures 3 and S1).

**Figure 2.**
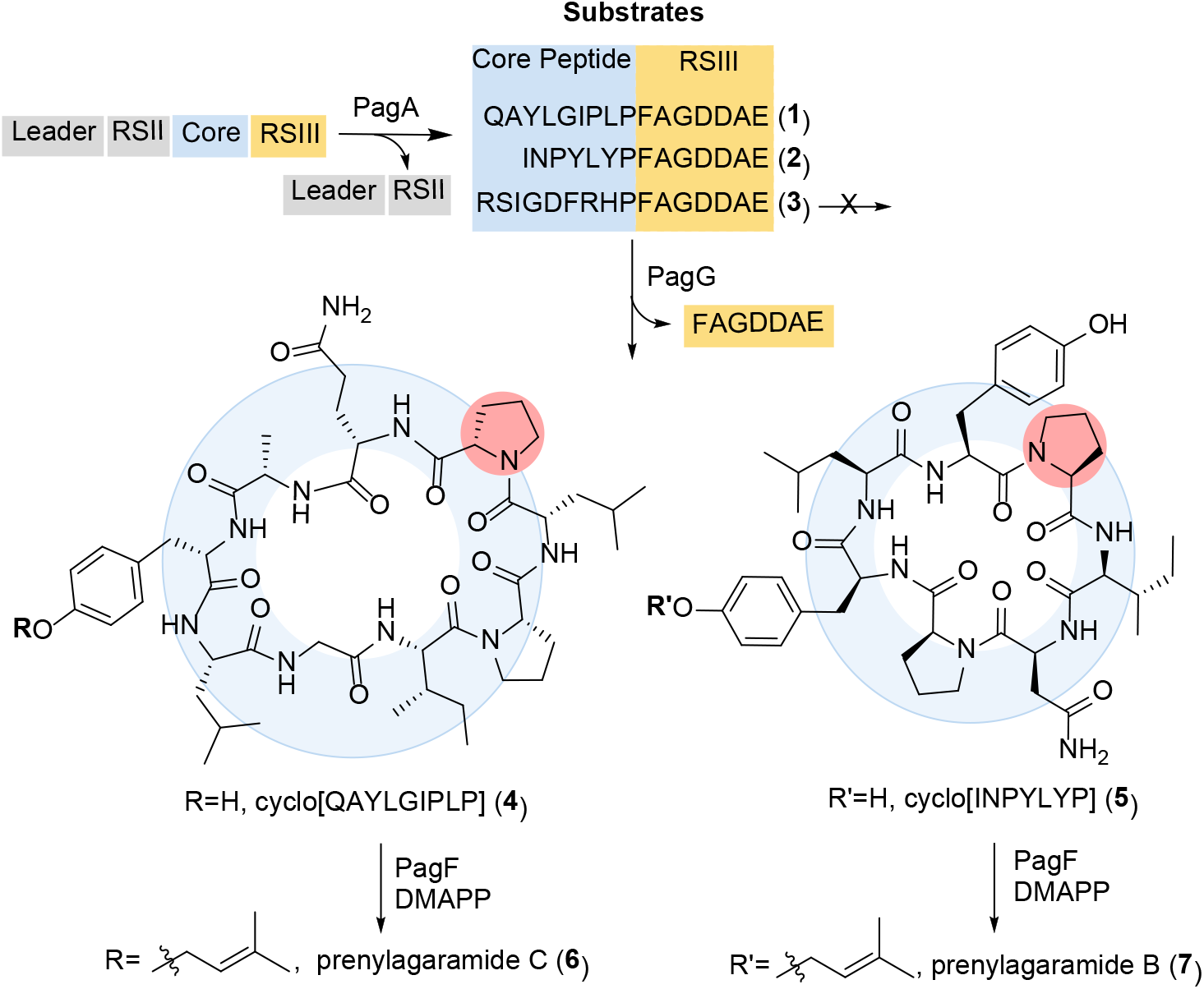
Proposed biogenesis of prenylagaramides. Precursor peptides are synthesized on the ribosome, and subsequently cleaved by PatA protease between RSII and the core peptide to release short substrates for macrocyclization (**1**-**3**). PatG protease cleaves the core peptide from RSIII (FAGDDAE), releasing the short peptide product and macrocyclizing the core peptides to yield products (**4**) and (**5**). Finally, the cyclic peptide is prenylated using DMAPP by enzyme PagF to produce the natural compounds, prenylagaramides C (**6**) and B (**7**).

**Figure 3.**
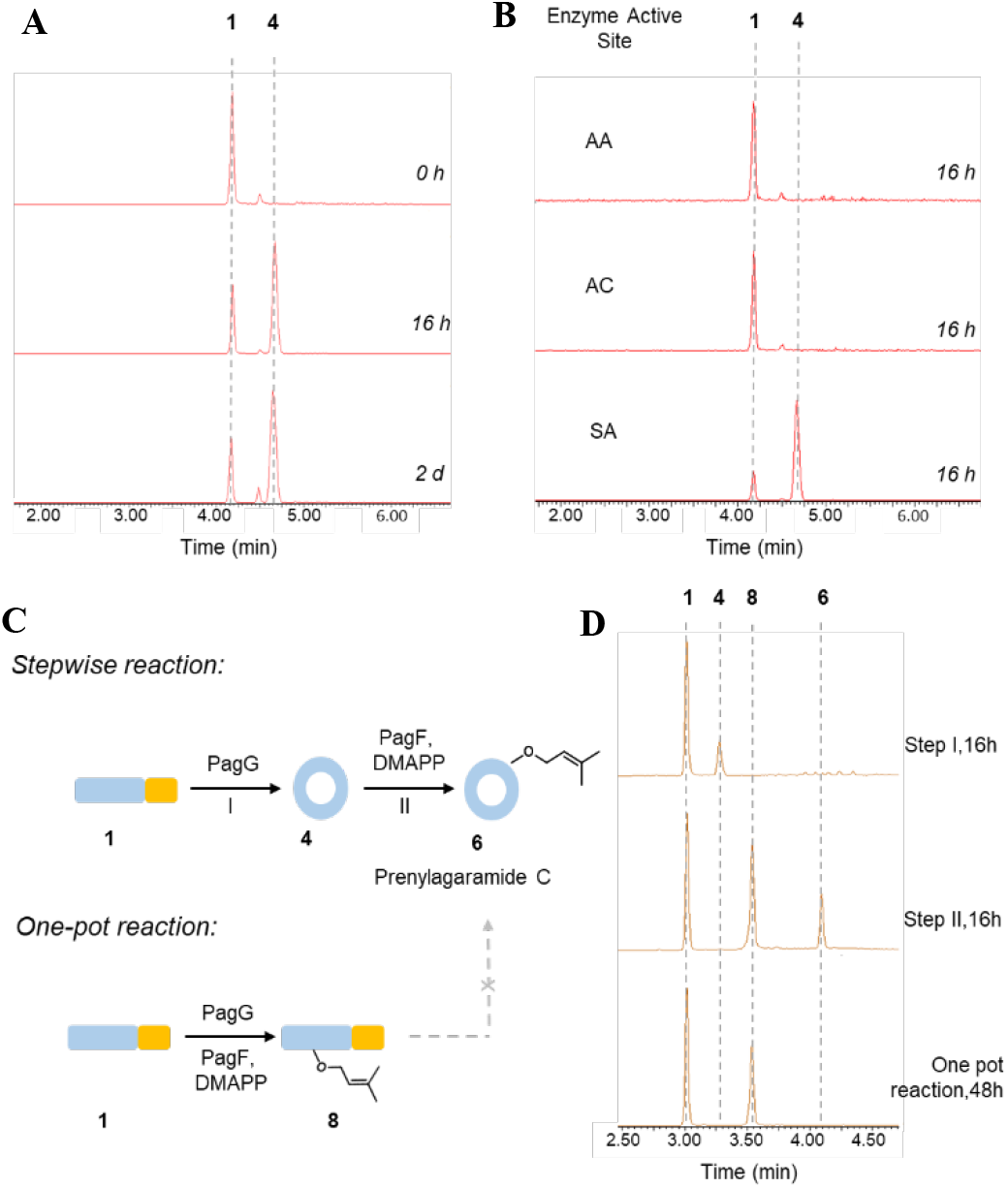
*in vitro* PagG activity. (A) Extracted ion chromatograms of substrate **1** and product **4** at 0, 16 and 48 h of *in vitro* assay generated through LC/MS. (B) Extracted ion chromatograms of substrate **1** and product **4** at 16 h of *in vitro* assay with S274A/C275A, C275A and S274A generated through LC/MS. (C) Synthetic scheme for stepwise and one-pot reactions. (D) Extracted ion chromatograms of the reactant, intermediates and product involved in the synthetic scheme.

In the canonical cyanobactin biosynthesis, a precursor peptide is cleaved after RSII, liberating an amino group that is the nucleophile for the subsequent macrocyclization event. Once RSII is cleaved, the remaining short precursor peptide should be the substrate for PagG^mac^. It contains only the core peptide, encoding the natural product, and RSIII, which is recognized by PagG^mac^. Therefore, we used synthetic native substrates (**1**-**3**) with PagG^mac^ (Figure 2A).

Two of these substrates, **1** and **2**, are natively converted into natural products (Figure 2), while the product of the third, **3**, has not been isolated even though the precursor peptide is found in the *P. aghardii* genome.^34^ When **1** or **2** were treated with PagG^mac^, cyclic products were **4** and **5** were obtained (Figure 3A), but substrate **3** was unreactive. Enzymatic reaction products **4** and **5** were compared with synthetic standards, confirming their structural identities (Figures S4-S7). These results showed that PagG^mac^ was active, and that its activity reflected expectation for the wild-type enzyme.

Like PatG, PagG^mac^ was a slow enzyme, with several hours required for significant product formation. Extensive reaction optimization was attempted, using different buffers, pH, temperature and additives (Figure S8), but velocity was not significantly improved. In presence of 1:1 substrate to enzyme ratio, a velocity of 370 nM/h was observed (Figure S9). We speculate that conditions do not replicate what is found in the cytoplasm of cyanobacteria, and specifically that a better understanding of the role of DUFs may improve yield.

### PagG^mac^ active site mutational analysis

PatG and related macrocyclases are subtilisin-like serine proteases, which have been extensively characterized via mutagenesis and structural studies. PagG^mac^ and its relatives share some similarities with PatG, including a similar predicted secondary structure and the Asp29-His113-Ser274 catalytic triad. However, there are several differences, including an unusual Cys275 in the active site adjacent to the nucleophilic Ser274 residue, while in PatG and relatives this residue is methionine (Figure S10). This is unique as both the serine and cysteine can act as a nucleophile for proteolysis, but cysteine has higher nucleophilicity due to the lower pKa of the sulfhydryl group.

The Ser274-Cys275 diad in the catalytic center was examined by simultaneously mutating both residues, to create seven point mutant proteins (Table 1, Figure S11). These mutants were assayed using linear substrate **2**, and using mass spectrometry to quantify macrocyclic product **5**. Product **5** was only observed when using mutants Cys275Ala, Cys275Ser, and Cys275Met. Thus, Ser274 is required, but Cys275 is not important for activity, ruling out the role of the sulfhydryl nucleophile in catalysis (Figures 3B and S12). Ser274 could not be substituted with Cys. This is similar to what has previously been found in studies of PatG: since sulfur is larger than oxygen, when the active-site Ser is replaced with Cys, a concomitant expansion of the active site is required.^25^ The Cys275Ala active site mutant showed a comparable activity of 390 nM/h.

**Table 1.**
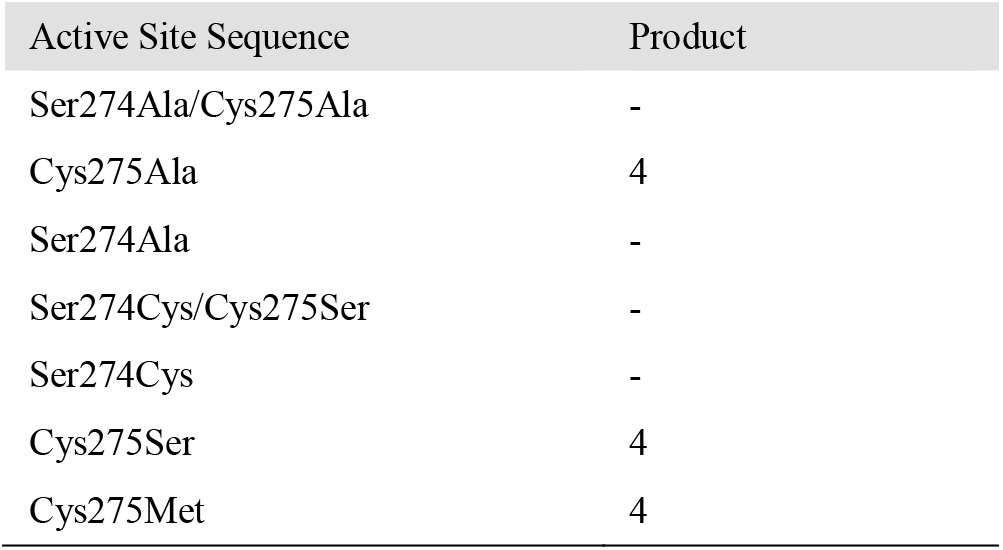
PagG^mac^ active site mutant reactions using substrate **1**

| Active Site Sequence | Product |
| --- | --- |
| Ser274Ala/Cys275Ala | - |
| Cys275Ala | 4 |
| Ser274Ala | - |
| Ser274Cys/Cys275Ser | - |
| Ser274Cys | - |
| Cys275Ser | 4 |
| Cys275Met | 4 |

### *in vitro* reconstitution of *pag* pathway and formal total synthesis of prenylagaramide B & C

*pag* is predicted to encode three enzymes: PagA, which proteolyzes the precursor peptide to release substrates **1**, and **2**, PagG, which macrocyclizes linear **1** and **2** to produce **4** and **5**, and PagF, which prenylates **4** and **5** to produce the natural products **6** and **7** (Figure 2A).^34^ Since we used synthetic short substrates **1** and **2**, *pag* would be reconstituted simply using PagG^mac^ and PagF; PagA is not required. In a stepwise procedure, treatment of substrate **1** first with PagG^mac^ and subsequently with PagF and dimethylallyl pyrophosphate (DMAPP) yielded the natural product **6**, representing the first total synthesis of **6**. In the second step in this procedure, the leftover substrate is also prenylated to form **8** (Figure 3).

A one-pot reaction containing all needed reagents failed because of the faster rate of PagF in comparison to PagG^mac^. Linear substrate **1** was prenylated prior to macrocyclization, yielding prenylated linear product **8**. We found that the linear peptides, once prenylated, are no longer substrates for macrocyclase (Figure 3). This defines the biosynthetic order of the *pag* pathway, demonstrating that the proposed route shown in Figure 2 is correct (Figures S13 and S14).

### Mutational analysis of the CH-RSIII interaction

Unlike canonical serine proteases, cyanobactin macrocyclases feature an additional CH, which binds to RSIII, enabling the macrocyclase to modify diverse core peptides.^25, 38^ This is a critical interaction that decouples the macrocyclase reaction from substrate recognition, making the enzymes ideal for biotechnology. The CH also stabilizes the tetrahedral orthoamide intermediates formed during the catalysis. Strikingly, when we compared the CH and RSIIIs from *pag* and related pathways, we observed entirely different sequence motifs, which further led to identification of a previously unremarked internal RS embedded *within* G-type cyanobactin macrocyclases (Figure 4 and S10).

**Figure 4.**
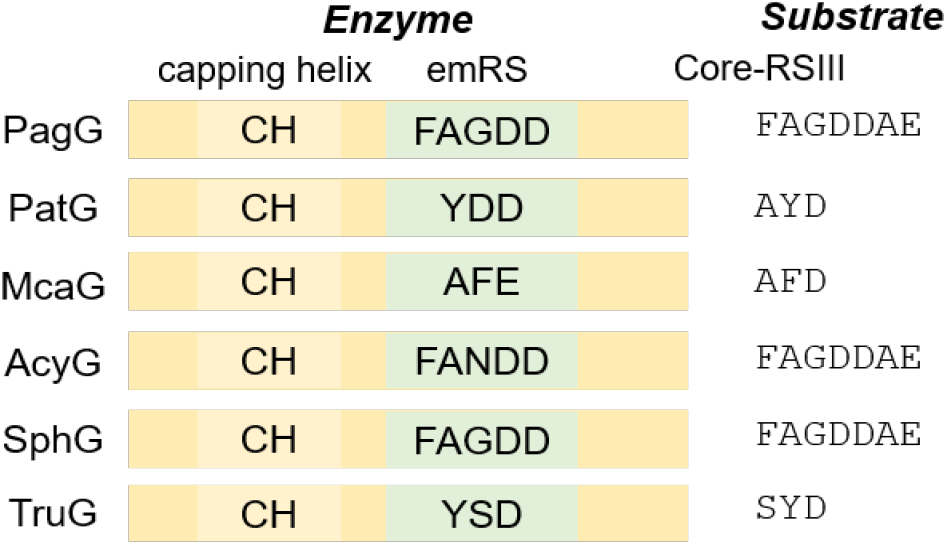
Embedded RS in cyanobactin G enzymes. In the G enzymes, an embedded RS (emRS) is present downstream of the capping helix (CH), which binds the substrate RSIII. The emRS in the enzyme closely resembles the predicted or characterized RSIII in the cognate substrates. We hypothesize that the CH binds emRS in the absence of substrate (Figure S15)

In PatG, structural studies revealed that CH is comprised of residues Pro579-Val605, which aligned with Asp59-Lys98 in PagG^mac^. While PatG binds to RSIII elements with sequences similar to AYDGE/SYDD, PagG^mac^’s RSIII sequence is FAGDDAE. A close examination of the G protein sequence alignment showed that an embedded recognition sequence (emRS) is present downstream of the CHs. These observations prompted us to hypothesize that the emRS might bind to the cap in the absence of the substrate. (Figure S15). In previous macrocyclase crystal structures, the emRS region was unstructured.^25, 38^ Hence, further insights could not be drawn from the published studies.

To test the hypothesis, we expressed a series of chimeric PagG^mac^ proteins wherein we swapped or added the PatG CH and emRS sequences (Table S8, Figure S16). Unfortunately, all constructs were soluble but inactive in our hands. In the D series of cyanobactin enzymes, recognition elements can be swapped using simple rules,^39^ but these rules are not as straightforward for the CH.

### Broad substrate tolerance of PagG^mac^

PatG and relatives are exceptionally substrate-tolerant enzymes.^29^ It was unclear whether other cyanobactin G enzymes would exhibit similar tolerance. Therefore, we performed saturation mutagenesis on each position P2-P7 of substrate **1**, leading to 114 unique peptide sequences and 6 duplicates of the native peptide sequence (Figure 5A). When treated with PagG^mac^, we found that ~60% of the substrates led to robust macrocyclization (Figure 5B). Only ~19% of the peptides remained unmodified by PagG^mac^. These results are similar to what as previously found with PatG, indicating that PagG^mac^ is a promising, broad-substrate catalyst. A few rules regarding the substrate tolerance of PagG^mac^ could be discerned: acidic residues (Asp/Glu) are not tolerated at any position; positions P4-P6 are more accepting of variation than P2, P3, or P7.

**Figure 5.**
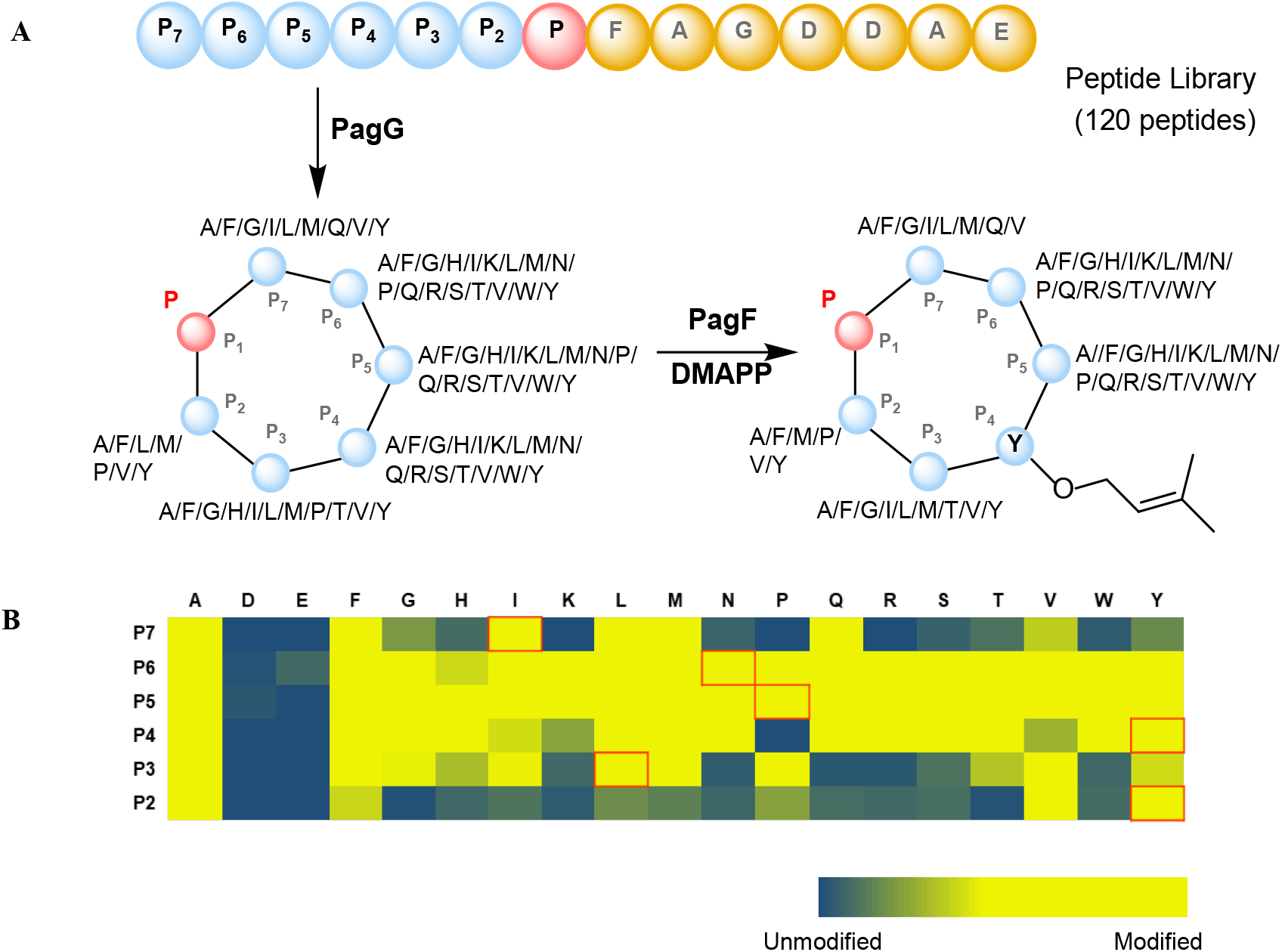
PagG^mac^ is a promiscuous catalyst. (A) A peptide library was synthesized in which each position P_2_ to P_7_ is independently mutated by each of the 20 proteinogenic amino acids. The library was arrayed into wells and treated with PagG^mac^, followed by PagF. The resulting macrocycles are shown schematically, with each accepted amino acid listed adjacent to each residue. (B) A heat map showing the acceptability of amino acids in positions P_2_ to P_7_ for macrocyclization by PagG^mac^. The heat map was generated using LC-MS traces for each reaction. Wild type residues are boxed in red.

Some of these rules make sense in light of the elegant PatG crystal structures. ^25, 38^ In the structure of the PatG complex with pentapeptide PIPFP, PatG causes the linear substrate to adopt a curved shape, bringing the N- and C-termini into proximity. In addition, the substrate P2 and P3 residues make limited van der Waals contact with the protein surface, while P4 and P5 make no contacts with PatG. Here, we observed that positions P4-P6 were more permissive, supporting the possibility that they have fewer contacts with PagG^mac^. We also propose that the inability to tolerate D/E in the core might be related to an incompatibility with the highly acidic RSIII sequence. Taken together, these rules will be useful in applying PagG^mac^ to synthetic biology or drug design problems.

To validate the library approach, we performed a time course study with three randomly chosen peptides from the library using both wild-type PagG^mac^ and the C275A active-site mutant. The reactions were complete in 14 and 10 hours for the wild-type and mutant proteins, respectively (Figure S18). One recurring disadvantage of the macrocyclases from cyanobactins is that they are relatively slow.

Here, we show that PagG^mac^ is faster than the previously characterized enzymes, being complete in 14 hours with a 20% catalyst load (Figure S18H), whereas PatG is complete after 24 hours using the same conditions.^30^ Further knowledge of internal mechanisms involving the DUF domain and capping helices may help to solve this problem

### Substrate tolerance of PagF and use in library generation

Although PagF was previously characterized with several substrates, its application to cyclic peptide libraries has not been probed.^37^ We used the macrocyclization reaction library as substrates for PagF prenylation (Figures 5A and S17). Each well in the library contained the unreacted linear substrate peptides, as well as cyclic products where they could be formed as shown in Figure 5A. PagF modified virtually every peptide, as long as Tyr is present at P4 (the natively prenylated residue). Both cyclic and linear peptides were prenylated on Tyr at P4 position. Thus, PagF is also tolerant of significant sequence variation. By combining PagG^mac^ and PagF, we synthesized a complex and chemically rich library of peptides modified by natural posttranslational enzymes.

### PagG^mac^ represents the first characterized representative of a new class of macrocyclases

Only a few of the cyanobactin cyclases have been previously characterized.^23, 26, 34, 40–41^ We obtained 65 cyanobactin macrocyclase and related protein sequences, which are broadly representative of all cyanobactin G-proteins, and performed a Bayesian phylogenetic analysis (Figure 6). For many of the G protein sequences, we could also find substrate precursor peptides in the adjacent gene clusters and assign their functions based upon known natural products. Strikingly, these proteins grouped into clades that are based upon their natural substrate selectivity.^33–35, 42–50^ Our analysis thus identifies a series of catalysts that may be used to macrocyclize different types of peptides or peptide libraries. Among these, only four (including PagG^mac^) have been biochemically characterized, and only two (TruG and PagG^mac^) have been well characterized with defined substrate libraries. However, for many of the uncharacterized proteins, the known substrates and products are quite variable, indicating that broad substrate tolerance is a feature of cyanobactin G proteins.

**Figure 6.**
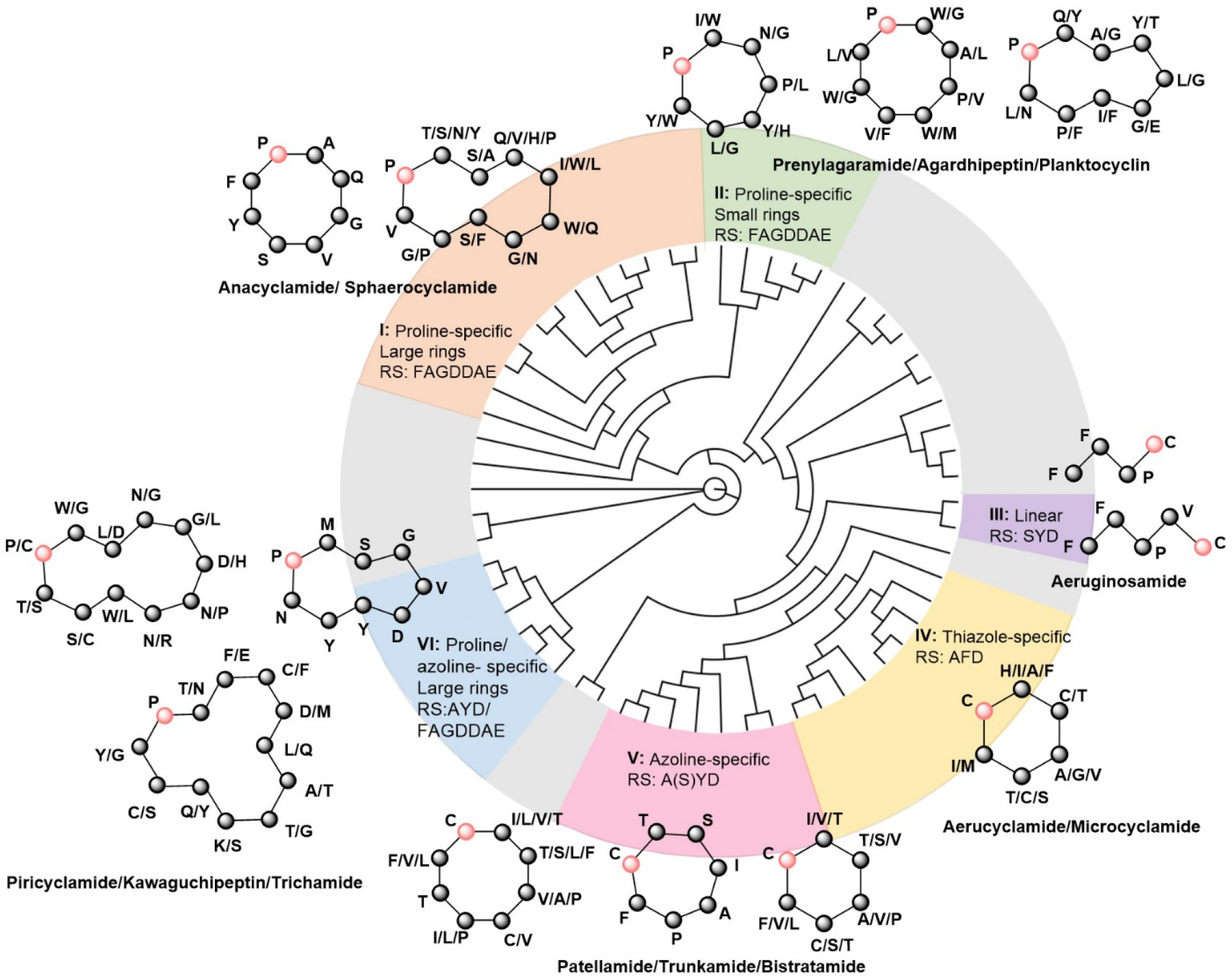
Native chemical space of cyanobactin macrocyclases. Bayesian phylogenetic tree of 65 cyanobactin C-terminal protease/macrocyclases. For each of six color-coded enzyme classes, known product structures are shown. The C-terminal macrocyclization site in each product is indicated with a red bubble. Close observation shows that for all of the substrate selectivity “rules” observed for PagG^mac^ or PatG / TruG, there are existing macrocyclases that break those rules and have complementary selectivity. Thus, judicious macrocyclase selection enables synthesis of diverse cycles. See Figure S19 for a complete tree with names and bootstrap values.

Phylogenetic analysis demonstrates that *any* gap in the substrate tolerance of PagG^mac^ or TruG can be filled by the other classes (Figure 6). For example, PagG^mac^ does not accept glycine, serine, or threonine at position P2, but those residues are found in the natural products of proline-specific macrocyclases from class VI. Peptides with proline at P4 position can be cyclized by class II macrocyclases (Figure 5B). Alternatively, TruG disfavors histidine at P3 or tyrosine at P4, but PagG^mac^ processes such substrates (Figure S20).^29^ While acidic amino acids were not accepted by PagG^mac^, class II cyclase products contain those residues. Thus, for each limitation of PagG^mac^ or TruG/PatG, there exists a cyclase that circumvents that limitation.

Moreover, several clades have no characterized representatives, and we cannot predict their substrates or products from available data. These represent promising areas for further biochemical research. Combined with macrocyclases from other classes of biosynthesis, there is a growing arsenal of enzymes to tackle challenging problems in macrocycle synthesis. The complete phylogenetic analysis is given in the supporting information (Figure S19).

A frontier of cyanobactin macrocyclases involves their demonstrated ability to circularize substrates that diverge significantly from ribosomal peptides, for example containing D-amino acids, polyketide-like spacers, and other nonproteinogenic components.^26–27^ The substrate scope for non-protein cyclization cannot be discerned from the phylogenetic tree and requires experimental evidence.

## CONCLUSION

We present the biochemical characterization and substrate tolerance profile of a new class of macrocyclase enzymes. We characterized the prenylagaramide biosynthetic pathway and synthesized natural products for the first time. Additionally, we used the tools from this pathway to generate a library of macrocyclic and prenylated peptides, demonstrating promising broad-substrate tolerance. We show that the natural cyclases group by substrate preference. Because of the many different macrocyclization catalysts available in this family, and the clear substrate rules delineated above, we predict that judicious choice of enzyme and substrate will enable the macrocyclization of virtually any linear compound.

## ASSOCIATED CONTENT

### Supporting information

The Supporting Information is available free of charge on the ACS Publications website at DOI:

Experimental methods, additional figures and tables (PDF)

## AUTHOR INFORMATION

### Notes

The authors declare no competing financial interests.

## ACKNOWLEDGMENTS

This work was funded by NIH GM122521. S.S. is supported by the Skaggs Graduate Research Fellowship. We thank Elizabeth Pierce and Maho Morita for preparing the PagF enzyme and DMAPP substrate used in this study.

